# The combined use of scRNA-seq and network propagation highlights key features of pan-cancer Tumor-Infiltrating T cells

**DOI:** 10.1101/2024.10.31.621221

**Authors:** Adèle Mangelinck, Elodie Molitor, Ibtissam Marchiq, Lamine Alaoui, Matthieu Bouaziz, Renan Andrade-Pereira, Hélène Darville, Etienne Becht, Céline Lefebvre

## Abstract

Improving the selectivity and effectiveness of drugs represents a crucial issue for future therapeutic developments in immuno-oncology. Traditional bulk transcriptomics faces limitations in this context for the early phase of target discovery as resulting gene expression levels represent the average measure from multiple cell populations. Alternatively, single cell RNA sequencing can dive into unique cell populations transcriptome, facilitating the identification of specific targets. Here, we generated Tumor-Infiltrating regulatory T cells (TI-Tregs) and exhausted T cells (Tex) gene signatures from a single cell RNA-seq pan-cancer T cell atlas. To overcome noise and sparsity inherent to single cell transcriptomics, we then propagated the gene signatures by diffusion in a protein-protein interaction network using the Patrimony high-throughput computing platform. This methodology enabled the refining of signatures by rescoring genes based on their biological connectivity and shed light not only on processes characteristics of TI-Treg and Tex development and functions but also on their immunometabolic specificities. The combined use of single cell transcriptomics and network propagation may thus represent an innovative and effective methodology for the characterization of cell populations of interest and eventually the development of new therapeutic strategies in immuno-oncology.

## Introduction

Since the first report of single cell transcriptomics in 2009 (1), this powerful approach has known a tremendous technological and computational development. It is now possible to sequence tens of thousands of cells in parallel and integrate readouts from DNA, RNA and protein with multi-modal single cell methodologies (2, 3). This enables cellular heterogeneity and interactions to be dissected more precisely than ever before. As such, single cell RNA sequencing (scRNA-seq) has led to ground-breaking discoveries in several fields including development, oncology, immunology, and neuroscience.

However, scRNA-seq protocols face technical limitations. One that feeds the majority of computational challenges is the low mRNA capture efficiency, leading to so-called dropout events (4). These dropout events consist in genes displaying null expression levels in cells (since none of their transcripts may be captured) while being observed at low or moderate expression levels in other cells of the same type. This issue causes both noise and sparsity of data (5). A common practice to address these dropout events is to preprocess the data by selecting highly variable genes to perform dimension reduction. Imputation and denoising methods have also been developed to estimate correct expression levels in each single cell (6). These methods usually rely on various means of local averaging or regression but recent cell annotation tools like netImpute and netSmooth alternatively proposed to use network propagation for scRNA-seq data imputation (7, 8). They rely on the principle that genes related to a phenotype are more likely to have biological interactions with each other than with randomly chosen genes. Network propagation has also been used for various purposes in biology, including gene function prediction, disease characterization and drug target prediction (9).

Servier built up a high-throughput computing platform named Patrimony with the aim of capitalizing on both proprietary and public data to foster drug discovery (10). This platform integrates a large variety of data including a protein-protein interaction (PPI) network and a propagation or diffusion algorithm. It has been previously applied to target prioritization in immune diseases (such as primary Sjögren syndrome and systemic lupus erythematosus) and neurodegenerative diseases (including amyotrophic lateral sclerosis and Parkinson’s disease), drug repurposing in SARS-CoV-2 and pediatric cancers as well as identifying drug combinations in autoimmune diseases (11–13).

In this study, we created a scRNA-seq pan-cancer T cell atlas comprising published data from 249,587 cells across 159 samples in total. We then generated gene signatures for Tumor-Infiltrating regulatory T cells (TI-Tregs) and exhausted T cells (Tex) from this atlas. Subsequently, we identified key genes and molecular mechanisms associated with each signature by combining them with Patrimony using network propagation.

## Materials and Methods

### Data collection, preprocessing and T cell identification

The scRNA-seq data were collected from previously published datasets, adhering to the following selection criteria: 1) presence of T cells, 2) treatment-naïve patients, 3) solid tumors, and 4) inclusion of at least tumor and blood samples.

Each scRNA-seq dataset underwent separate preprocessing in R (v4.0.2). We filtered out cells from the original count matrices that had fewer than 200 genes detected or more than 10% mitochondrial UMI counts and we only kept genes detected in at least 3 cells. Then, we applied Seurat (v4.0.5) with default parameters for count data normalization and scaling. Each cell was assigned a cell cycle score using the *CellCycleScoring* function and we computed the difference between the G2M and S phase scores. This approach allows for the separation of non-cycling from cycling cells while minimizing the differences in cell cycle phase among proliferating cells. The *SelectIntegrationFeatures* function was ran with the nfeatures parameter set to 3,000 before merging all samples from each dataset. These integration features were then used for Principal Component Analysis (PCA) and Uniform Manifold Approximation and Projection (UMAP). Clustering was performed using the Louvain algorithm with the resolution parameter set to 2.0 for all datasets. Finally, T cells were isolated based on CD3D and CD3G genes expression (CD3D or CD3G expression level > 0).

### Construction of the T cell atlas

To integrate heterogeneous data from different sources, a two-step procedure was applied. We first concatenated all datasets together and ran the scaling and PCA steps based on the top 3,000 highly variable genes identified by the *FindVariableFeatures* function with the “vst” method. Harmony was applied for batch effect correction then UMAP and clustering using the Louvain algorithm with the resolution parameter set to 2.0 were performed on the harmony reduction. Examining the result from the first clustering run, we identified contamination clusters and clusters that arose from unwanted factors: we removed the contamination clusters including low quality cells highly expressing marker genes associated with apoptosis and tissue dissociation operation, pancreatic acinar cells (expressing PRSS1, CLPS, PNLIP and CTRB1 among others), myeloid cells (expressing CD68) and B cells (expressing CD79A). Then, we performed the second run of integration and clustering excluding immunoglobulin, ribosome-protein-coding, and T cell receptor (TCR) genes (gene symbol with string pattern “^IGK|^IGH|^IGL|^IGJ|^IGS|^IGD|IGFN1”, “^RP([0-9]+-|[LS])”, and “^TRA|^TRB|^TRG” respectively) from the top 3,000 highly variable genes and regressing out the cell cycle difference effect as well as the percentage of mitochondrial UMI counts. Harmony (14) (v0.1.0) was applied again for batch effect correction and UMAP was performed on the harmony reduction.

T cell subtypes identification and annotation was performed by clustering cells using the Louvain algorithm with the resolution parameter set to 4.1 after iterative testing from 3.5 to 5.0 by 0.1 (more granular than default), computing clusters signatures based on differential gene expression using the *FindAllMarkers* function with the “MAST” method and interrogating known gene markers expression.

Figures were generated in R (v4.3.1) using the tidyverse (v2.0.0), ggpubr (v0.6.0), ComplexHeatmap (v2.16.0) (15), circlize (v0.4.15) (16) and viridis (v0.6.4) packages.

### Generation of gene signatures

T cell subtypes gene signatures were generated in R (v4.3.1) using either the *FindAllMarkers* or the *FindMarkers* function from Seurat (v4.4.0) with the batch effect variable specified in the latent.vars, *MAST* method and default thresholds. We further selected genes exhibiting a positive log2 fold change and an adjusted p-value inferior to 0.05 (based on Bonferroni correction).

The *FindAllMarkers* function was used to conduct differential expression analyses (1) between the Treg population and all other CD4^+^ T cells or (2) between the Tex population and all other CD8^+^ T cells within intra-tumor CD4^+^/CD8^+^ subsets of the pan-cancer scRNA-seq T cell atlas, defining respectively the “Treg vs all” and “Tex vs all” signatures.

The *FindMarkers* function was used to conduct differential expression analyses (1) between the Treg and Thelper populations or (2) between the Tex and Tcyt populations within intra-tumor CD4^+^/CD8^+^ subsets of the pan-cancer scRNA-seq T cell atlas, defining respectively the “Treg vs Thelper” and “Tex vs Tcyt” signatures.

### Network propagation analysis

Network propagation analysis was performed on a high-throughput computing platform named Patrimony (10, 13).

Patrimony relies on a human PPI network and the diffusion of a score in this network. The PPI network was derived from STRING (v11.5) (17) and RNAinter (v4.0) (18, 19) databases. Links with experimental evidence and a *combined_score* superior to 300 were selected from STRING while links from human-derived data with a *confidence_score* superior to 0.25 from RNAinter were included. As such, the PPI network in Patrimony actually comprises both protein-protein interactions and RNA-protein interactions all annotated at the gene level but the term of PPI network was kept for simplicity matters.

We assigned to each protein in the network a *biological relevance score*: the log2 fold change value from the differential expression analysis for proteins encoded by genes in the scRNA-seq signatures and 0 for all other proteins.

Then, the algorithm in Patrimony is a lazy random walk using the heat kernel method (20, 21). Its implementation was based on Cowen and co-workers’ (9), using the *laplacian* function from the python library “graph_tool” (https://graph-tool.skewed.de/) and the *linalg.expm* function from scipy library (https://docs.scipy.org/doc/scipy/reference/generated/scipy.linalg.expm.html).

First the adjacency matrix A and the matrix D of the network are calculated, with Graph G = (V, E) where V = ensemble of n vertices & E = ensemble of edges

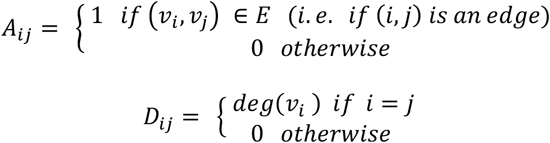

Then the Laplacian matrix L = D – A is calculated:

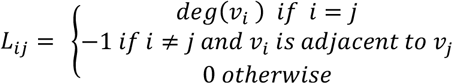

Finally, to estimate how information is propagated through the network, the similarity matrix W is calculated as follow: *W* = e^−*αL =*^ e^−*α*(*D* − *A*)^ with α being a decay factor of the diffusion and equal to 0.1 in practice (22).

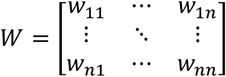

The W matrix can be calculated only for undirected networks (weighted or not), but not for directed ones. Indeed, the Laplacian matrix L in that case is symmetric, which is required to apply diffusion kernel. Indeed, if the graph is directed, then the adjacency matrix A is asymmetric, and so is the Laplacian matrix L.

Then, a *diffusion score* is calculated using the previously calculated scores (in this study, the *biological relevance score*) as follows, with S a vector of one of these scores and P the resulting vector after diffusion of these scores:

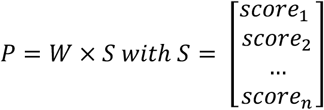

The *diffusion score* for gene g is:

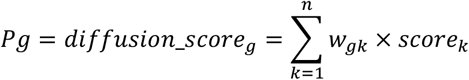

Of note, this diffusion algorithm was shown to more accurately represent distance in a PPI network than topological distance as it considers the local connectivity (20).

Figures were then generated in R (v4.3.1) using the igraph (v1.5.1) package with the Fruchterman-Reingold layout algorithm (force-directed placement of nodes).

### Functional analysis

Based on differentially expressed genes (DEGs) identified from scRNA-seq data or Patrimony-computed top gene signatures, Reactome analysis was performed using the https://maayanlab.cloud/Enrichr/ website (23–25). Figures were then generated in R (v4.3.1) using the tidyverse (v2.0.0) package.

### Statistical analysis

Statistical analysis was conducted in R (v4.3.1). One-way analysis of variance (ANOVA) with Tukey’s multiple comparisons tests were performed using the multcomp package (v1.4-25) for multiple group comparisons. For all tests, a p-value inferior to 0.05 was considered statistically significant.

## Data Availability

The data analyzed in this study were obtained from Gene Expression Omnibus (GEO) at GSE140228, GSE139555, GSE155698, GSE121636, GSE139324. The single cell RNA-seq pan-cancer T cell atlas generated from these datasets has been deposited in zenodo database under accession number 10.5281/zenodo.13879752.

## Results

### Construction of a pan-cancer single cell RNA-seq atlas of T cells

To generate a pan-cancer atlas of T cells, we collected scRNA-seq datasets including multi-tissues and blood samples from treatment-naïve cancer patients. Specifically, we compiled data from colorectal cancer (CRC) (26), intrahepatic cholangiocarcinoma (CHOL) and hepatocellular carcinoma (HCC) (27), head and neck squamous cell carcinoma (HNSCC) (28), non-small cell lung cancer (NSCLC) (26), pancreatic ductal adenocarcinoma (PDAC) (29), renal cell carcinoma (RCC) (30, 26) and uterine corpus endometrial carcinoma (UCEC) (26). After stringent quality control filtering, this atlas comprised data for 249,587 T cells from 64 patients and 15 healthy donors (Fig 1A-B), derived from tumor, adjacent normal tissue and blood samples among others (Fig 1C-D). Following integration, cells separated into 54 clusters that could be assigned to previously described T cell subtypes based on their expression signatures (Fig 1E-H, Fig S1A-C).

**Fig 1.**
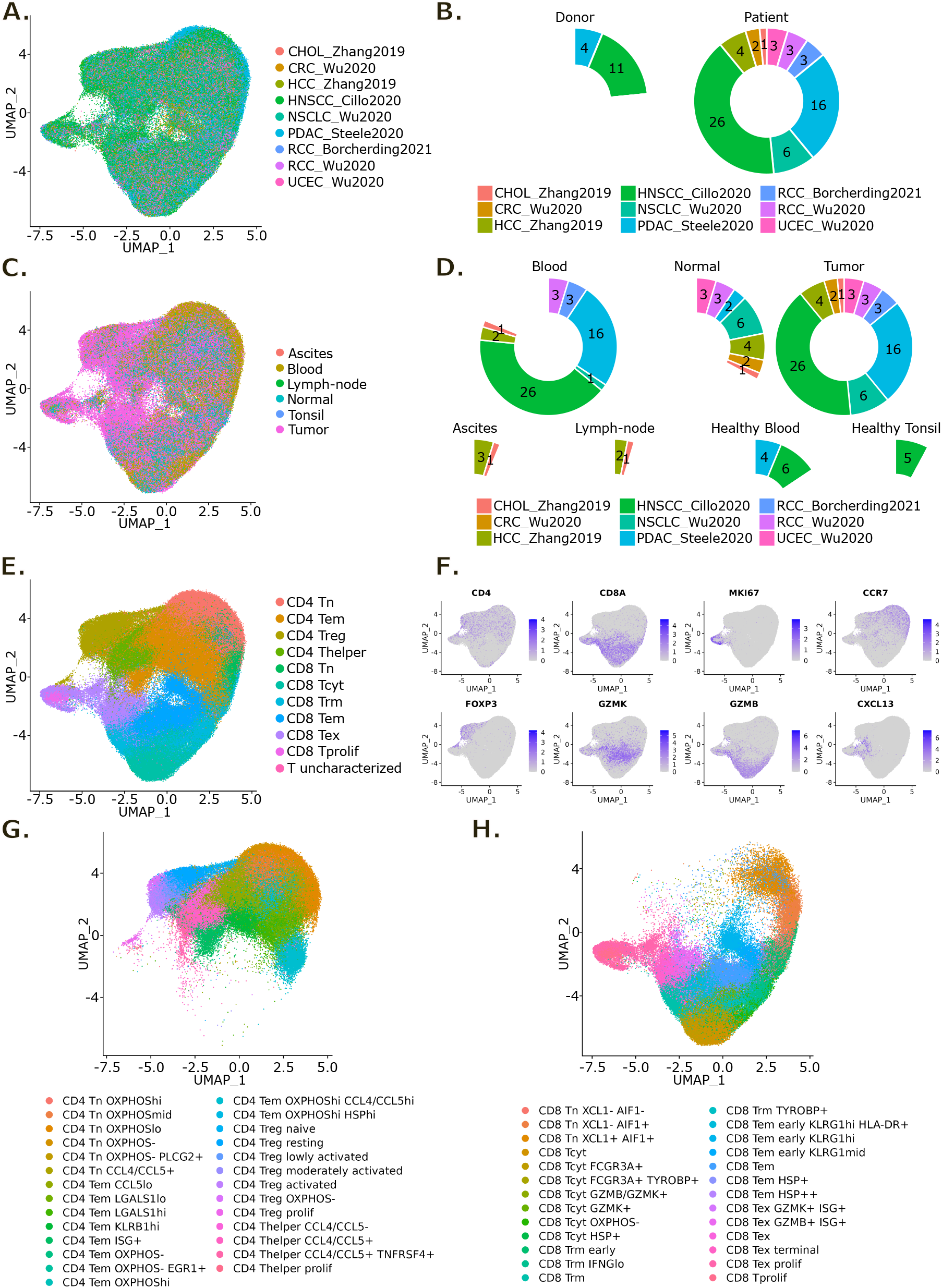
Construction of a pan-cancer scRNA-seq atlas of T cells. **(A)** UMAP representation of the T cell atlas colored by dataset. **(B)** Pie chart of the number of donors and patients per dataset. **(C)** UMAP representation of the T cell atlas colored by sample type. **(D)** Pie chart of the number of samples per sample type and dataset. **(E)** UMAP representation of the T cell atlas colored by main phenotype. **(F)** UMAP representation of RNA expression for the indicated genes in the T cell atlas. **(G, H)** UMAP representation of the T cell atlas colored by detailed CD4^+^ (G) and CD8^+^ (H) T cell phenotypes.

129,977 cells were identified as CD4^+^ T cells and 117,731 cells were identified as CD8^+^ T cells (Fig S1A). This represents a 52:48 ratio indicating close to equal proportions of these two T cell phenotypes. Among the CD4^+^ T cells, we identified 6 subtypes of naïve T cells (Tn) expressing TCF7, LEF1, CCR7, SELL and MAL that mainly separated on oxidative phosphorylation-related genes expression, 10 subtypes of effector memory T cells (Tem) that separated on both oxidative phosphorylation and activation-related genes expression (namely LGALS1, KLRB1 and interferon-stimulated genes) as well as 7 subtypes of regulatory T cells (Treg) expressing FOXP3, IL2RA and RTKN2, and 4 subtypes of T helper cells (Thelper) expressing TNFRSF18 but not Treg markers that separated on activation-related genes expression (mainly TNFRSF9, CCL4 and IL1R2 for Tregs, while CCL4 and CCL5 for Thelpers) (Fig 1G, Fig S1B). As for CD8^+^ T cells, we identified 3 subtypes of Tn expressing TCF7, LEF1, CCR7, SELL and MAL that separated on a few activation-related genes expression (namely XCL1 and AIF1), 6 subtypes of Tem expressing high levels of GZMK and low levels of GZMB that separated on both activation (KLRG1, HLA-DRA and NR4A1) and stress-related (HSPA1A, HSPA1B) genes expression, 7 subtypes of cytotoxic T cells (Tcyt) expressing GZMB, PRF1 and KLRG1 that separated on oxidative phosphorylation, activation (FCGR3A, TYROBP, GZMB/GZMK) and stress (HSPA1A, HSPA1B) -related genes expression as well as 4 subtypes of tissue-resident memory T cells (Trm) expressing ZNF683 and ITGAE and 5 subtypes of exhausted T cells (Tex) notably expressing CXCL13 and PDCD1 that separated on activation-related genes expression (mainly IL7R/GZMB and IFNG for Trm, while TNFRSF18 and ENTPD1 for Tex) (Fig 1H, Fig S1C). We could also identify a proliferating CD8^+^ population not clearly belonging to any of the other well-defined CD8^+^ T cell subtypes.

Cell composition analysis showed significantly higher frequencies of Treg and Thelper cells in tumors compared to both blood and normal tissues among the CD4^+^ T cell populations (Fig 2A-B). Treg proportions that represented 7.2 +/-2.6 % and 7.7 +/-2.6 % of CD4^+^ T cells respectively in blood and normal tissues reached 21.7 +/-11.4 % in tumors. As for Thelpers, their proportions increased from 4.2 +/-1.5 % in blood and 6.7 +/-4.0 % in normal tissue to 12.1 +/-6.3 % in tumors. Interestingly, this increase could be observed for all subtypes of Tregs and Thelpers, hence irrespectively of their activation level (Fig S2A). On the contrary, CD4^+^ Tn cells frequency was lower in tumors compared to blood and, along with Tem cells, acquired a phenotype associated with a metabolic switch toward oxidative phosphorylation (OXPHOS). As for CD8^+^ T cells, they featured high frequencies of Tex cells in tumors (22.3 +/-12.9 % of CD8^+^ T cells) and in particular terminally exhausted T cells, while this population was merely represented in blood and normal tissues (Fig 2C-D, Fig S2B).

**Fig 2.**
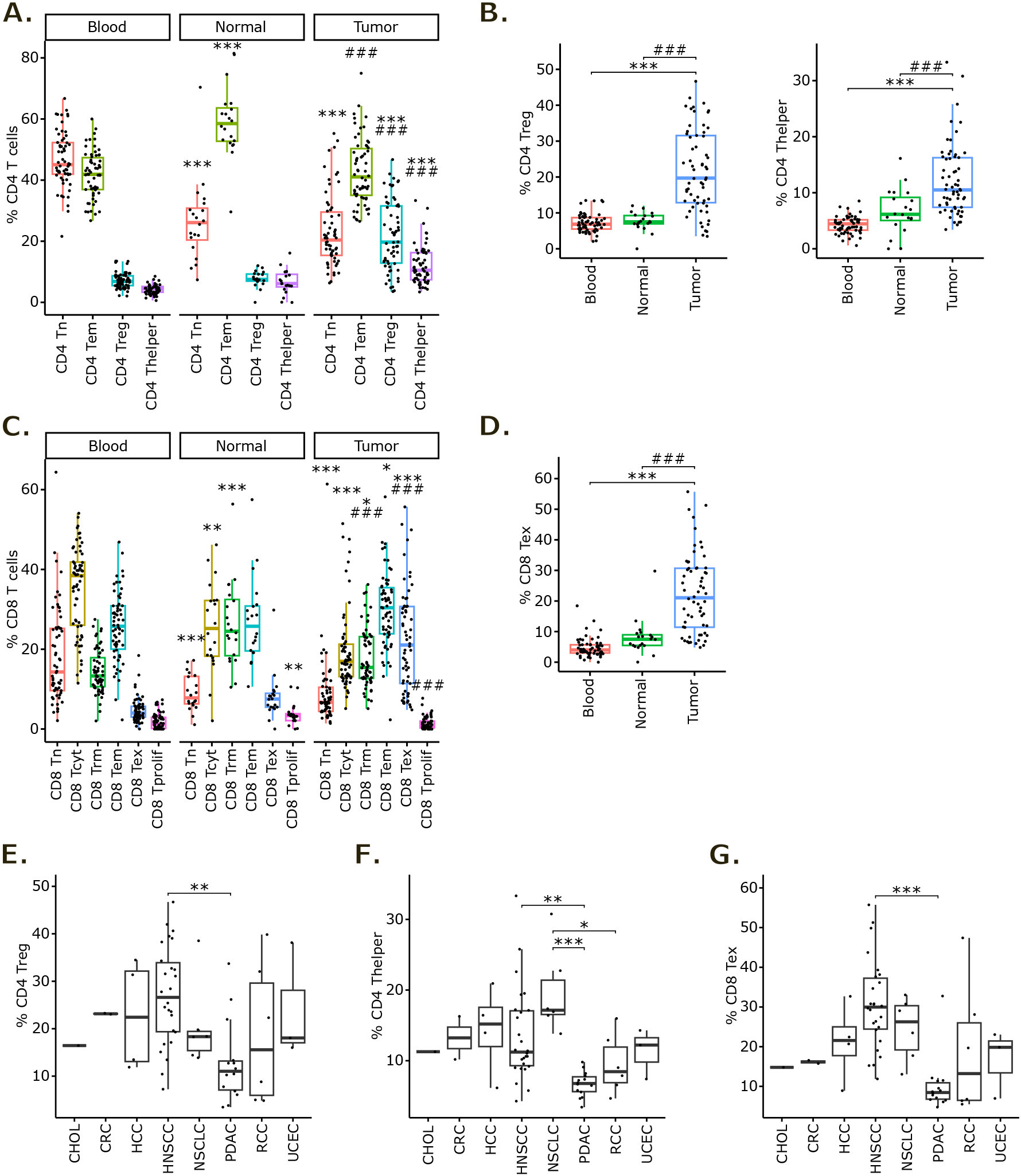
Cell composition analysis in the scRNA-seq T cell atlas. **(A)** Boxplot of CD4^+^ T cells main phenotype proportions compared to total CD4^+^ T cells in blood, adjacent normal tissue and tumor tissue. **(B)** Focused boxplots on Treg (left panel) and Thelper (right panel) populations. **(C)** Boxplot of CD8^+^ T cells main phenotype proportions compared to total CD8^+^ T cells in blood, adjacent normal tissue and tumor tissue. **(D)** Focused boxplot on Tex population. Statistical analyses for A to D: ANOVA with Tukey’s multiple comparisons tests. ^*^: p-value < 0.05, **: p-value < 0.01 and ***: p-value < 0.001 compared to blood sample proportion. #: p-value < 0.05, ##: p-value < 0.01 and ###: p-value < 0.001 compared to adjacent normal tissue sample proportion. **(E-G)** Boxplot of Treg (E), Thelper (F) and Tex (G) proportions compared to total CD4^+^ or CD8^+^ T cells in tumor samples across indications. Statistical analyses for E to G: ANOVA with Tukey’s multiple comparisons tests. *: p-value < 0.05, **: p-value < 0.01 and ***: p-value < 0.001 between indicated groups.

We then focused on intra-tumor Treg, Thelper and Tex proportions across cancer types and observed that these 3 populations’ proportions were lower in PDAC. This indicates that, compared to other solid tumors, T cell infiltrate in PDAC is less represented by Treg, Thelper and Tex cell populations (Fig 2E-G). PDAC is generally considered an immunologically “cold” cancer, characterized by low antigen presentation to the immune system, a dense and complex stromal microenvironment acting as a physical barrier, and an immunosuppressive milieu enriched with myeloid-derived suppressor cells (MDSCs) and M2 macrophages (31). These elements, in concert with the presence of immune checkpoint molecules and metabolic reprogramming within the tumor microenvironment, are key factors in T cell dysfunction and the resistance to immunotherapy. Collectively, these attributes likely account for the sparse presence of activated and immunotherapy-responsive T cell subsets in PDAC.

### Generation of a TI-Treg gene signature and its network-based rescoring

Here we aim to gain insight beyond scRNA-seq-derived gene signatures on pan-cancer TI-Treg markers. Therefore, to overcome the limitations of scRNA-seq drop out, we propose to combine TI-Treg signatures obtained from differential expression analysis with a network-based analysis. For this purpose, we first conducted a differential expression analysis between the Treg population and all other CD4^+^ T cells within an intra-tumor subset of the pan-cancer scRNA-seq T cell atlas, defining a “Treg vs all” signature. The up-regulated genes in TI-Tregs were enriched not only in genes specific to the Treg population such as FOXP3, IL2RA and RTKN2 but also genes more generally implicated in T cell activation like CD4, CD247, PTPN22, LCK and MHC class II genes (Table S1). Consequently, we also carried out a differential expression analysis between TI-Tregs and TI-Thelpers, defining a “Treg vs Thelper” signature, which successfully eliminated activation-related genes from this second Treg signature (Table S1). For both TI-Treg gene signatures, we computed a score for each intra-tumor CD4^+^ T cell of the atlas. UMAP representation of these scores showed that both TI-Treg gene signatures well characterized the intra-tumor Treg population for all datasets of the atlas (Fig. S2C). The two TI-Treg signatures, “Treg vs all” and “Treg vs Thelper”, shared 190 of their 571 and 240 up-regulated genes respectively (Fig 3A).

**Fig 3.**
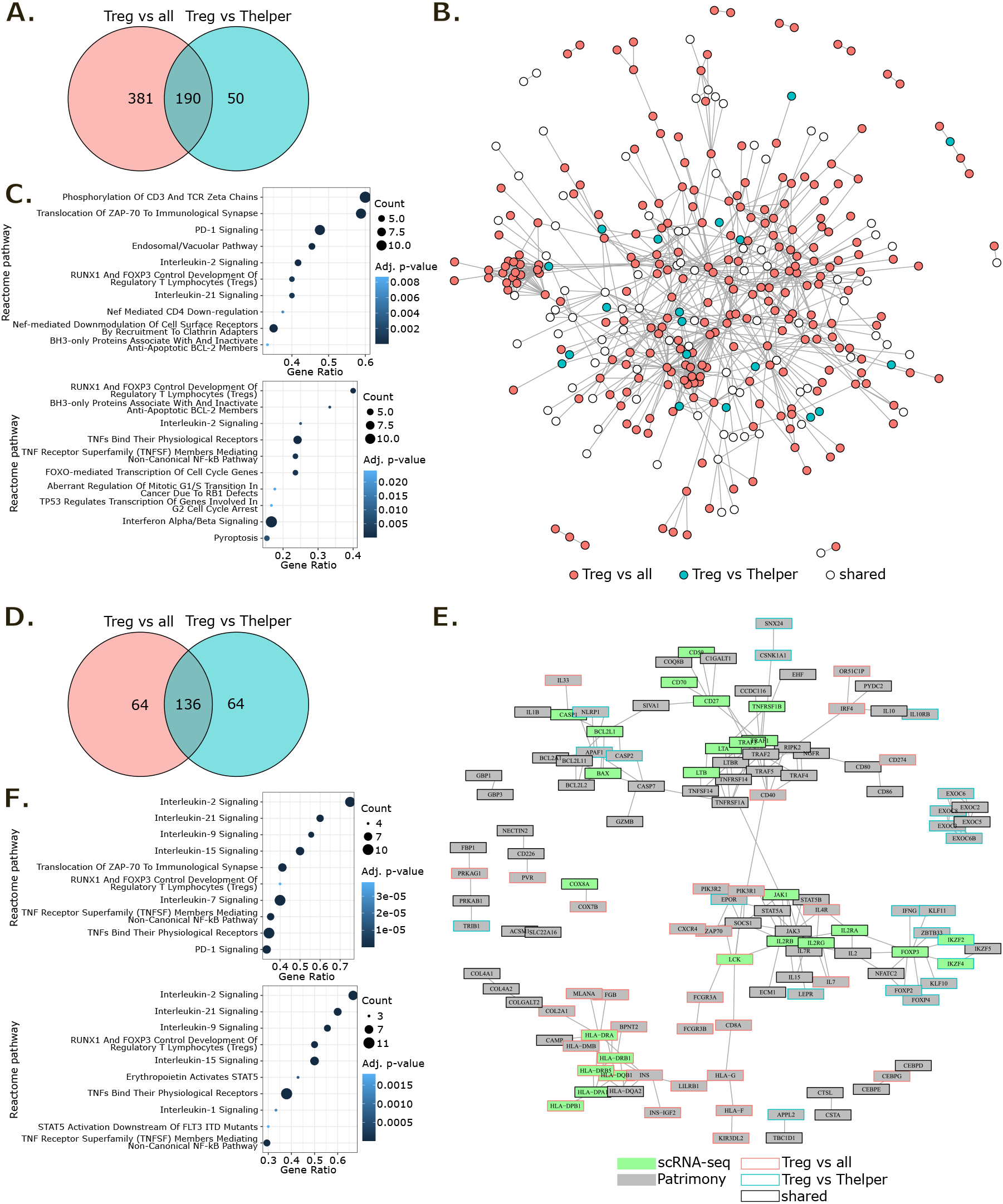
Generation of a TI-Treg signature from the T cell atlas and its network-based rescoring for target prioritization with Patrimony. **(A)** Venn diagram of TI-Treg scRNA-seq signatures. **(B)** Network representing direct interactions at the protein level between genes of TI-Treg scRNA-seq signatures. Genes with no direct interactions are not represented. **(C)** Top 10 enriched Reactome pathways in “Treg vs all” (upper panel) and “Treg vs Thelper” (lower panel) scRNA-seq gene lists. Pathways for which adjusted p-value < 0.05 and count > 2 were kept then ordered by decreasing gene ratio. **(D)** Venn diagram of TI-Treg Patrimony top 200 genes. **(E)** Network representing direct interactions at the protein level between genes of TI-Treg Patrimony top 200 results. Genes with no direct interactions are not represented. **(F)** Top 10 enriched Reactome pathways in “Treg vs all” (upper panel) and “Treg vs Thelper” (lower panel) Patrimony top 200 genes. Pathways for which adjusted p-value < 0.05 and count > 2 were kept then ordered by decreasing gene ratio.

The mapping of these signatures on Patrimony human PPI network revealed a high inter-connectivity between genes of the signatures that did not organize into distinct clusters (Fig 3B). Therefore, we conducted a pathway enrichment analysis to further investigate the scRNA-seq-derived TI-Treg signatures (Fig 3C). Pathways such as “RUNX1 and FOXP3 control development of regulatory T lymphocytes (Tregs)”, “BH3-only proteins associate with and inactivate anti-apoptotic BCL-2 members”, and “Interleukin-2 signaling” were among the top 10 enriched Reactome pathways for both TI-Treg signatures. The “Treg vs all” signature was also enriched in pathways related to TCR-activation and endosomal/vacuolar trafficking, while the “Treg vs Thelper” signature showed enrichment in pathways related to TNF signaling and cell proliferation.

The pathways enriched in the scRNA-seq TI-Treg signatures underscore well-documented molecular mechanisms critical for the development, survival and proliferation of Tregs, which involve the expression of FoxP3 and the modulation of anti-apoptotic Bcl-2 family members (32, 33). They also highlight enhanced immunosuppressive functions that have been described as dependent on sustained TCR and IL2 signaling (34). TI-Tregs display unique biological characteristics, making them more potent in fostering tumor progression. The enrichments observed in the endosomal/vacuolar trafficking pathway and the cell proliferation pathway within the “Treg vs all” and “Treg vs Thelper” signatures suggest that TI-Tregs have the capacity to modulate their metabolic processes to support both their immunosuppressive functions and proliferation within tumors (35).

We then performed computational network analysis using the Patrimony platform (10, 13) on the two TI-Treg scRNA-seq signatures. Briefly, a diffusion algorithm was applied to each of the two TI-Treg scRNA-seq signatures within the human PPI network. Consequently, every gene in the network (*i.e*. 18,787 genes) was assigned a diffusion score reflecting its connectivity to the input signature. Upon examining the distribution of scaled diffusion scores, we noted a rapid decrease from the top 1 to approximatively the top 200 scored gene for both TI-Treg Patrimony results (Fig S3A), suggesting that the diffusion process emphasizes a restricted number of genes with high connectivity to the input signatures. The “Treg vs all” and “Treg vs Thelper” Patrimony top 200 shared only two-thirds of their genes (Fig 3D, Table S1). Hence, even after Patrimony rescoring, the two signatures remained complementary. The mapping of Patrimony top 200 genes on the human PPI network revealed several gene clusters: a MHC class II genes cluster and a smaller one with MHC class I genes, a BCL2 family genes cluster, a cluster of exocyst-coding genes, as well as clusters centered around FOXP3, IL2 receptor genes, and TNF receptor genes (Fig 3E).

We also conducted pathway enrichment analysis on each TI-Treg Patrimony top 200 gene list (Fig 3F). Several interleukin signaling pathways (specifically, interleukin-2, -21, -9 and -15) were among the top 10 enriched Reactome pathways for both TI-Treg Patrimony top 200 gene lists. Additionally, the “RUNX1 and FOXP3 control development of regulatory T lymphocytes (Tregs)” and two TNF-related pathways: “TNF receptor superfamily (TNFSF) members mediating non-canonical NF-kB pathway” and “TNFs bind their physiological receptors” were also highly represented. The “Treg vs all” signature showed enrichment in the “Translocation of ZAP-70 to immunological synapse” and “PD-1 signaling” pathways. These two pathways were mainly driven by MHC class II genes. Conversely, the “Treg vs Thelper” signature was distinguished by an enrichment in STAT5-related signaling pathways.

TI-Treg signatures, both before and after network propagation, exhibit enrichment in closely related pathways. Therefore, diffusion in the PPI network did not fundamentally change the phenotypic characterization represented in the gene signatures. However, it more efficiently highlighted the complex interplay of signaling events that underpin the development, activation and maintenance of Tregs in the tumor microenvironment (TME). This methodology shed light on the role of Treg engagement in a multifaceted network of interactions within the TME, influencing immune responses through diverse mechanisms involving antigen processing and presentation, cell survival, cytokine signaling, and inflammation.

### Generation of a Tex gene signature and its network-based rescoring

We then applied our methodology to Tex marker identification. Two distinct gene signatures, “Tex vs all” and “Tex vs Tcyt”, were generated by conducting differential expression analyses between the Tex population and all other CD8^+^ T cells or the Tcyt population within an intra-tumor subset of the pan-cancer scRNA-seq T cell atlas (Table S1). For both Tex gene signatures, we computed a score for each intra-tumor CD8^+^ T cell of the atlas. UMAP representation of these scores showed that both Tex gene signatures well characterized the Tex population for all datasets of the atlas (Fig. S2D). The two Tex signatures showed a substantial overlap, with 549 shared up-regulated genes out of 614 and 633 respectively (Fig 4A). Similar to the TI-Treg findings, the mapping of these signatures onto the Patrimony human PPI network revealed a high inter-connectivity between genes of the signatures that did not organize into distinct clusters (Fig 4B). Consequently, pathway enrichment analysis was performed to discern the functional implications of the scRNA-seq-derived Tex signatures (Fig 4C). Pathways including the “Unwinding of DNA”, “Translocation of ZAP-70 to immunological synapse”, SLBP dependent processing of replication-dependent histone pre-mRNAs”, “Phosphorylation of CD3 and TCR zeta chains” and “PD-1 signaling” were among the top 10 enriched Reactome pathways for both Tex signatures.

**Fig 4.**
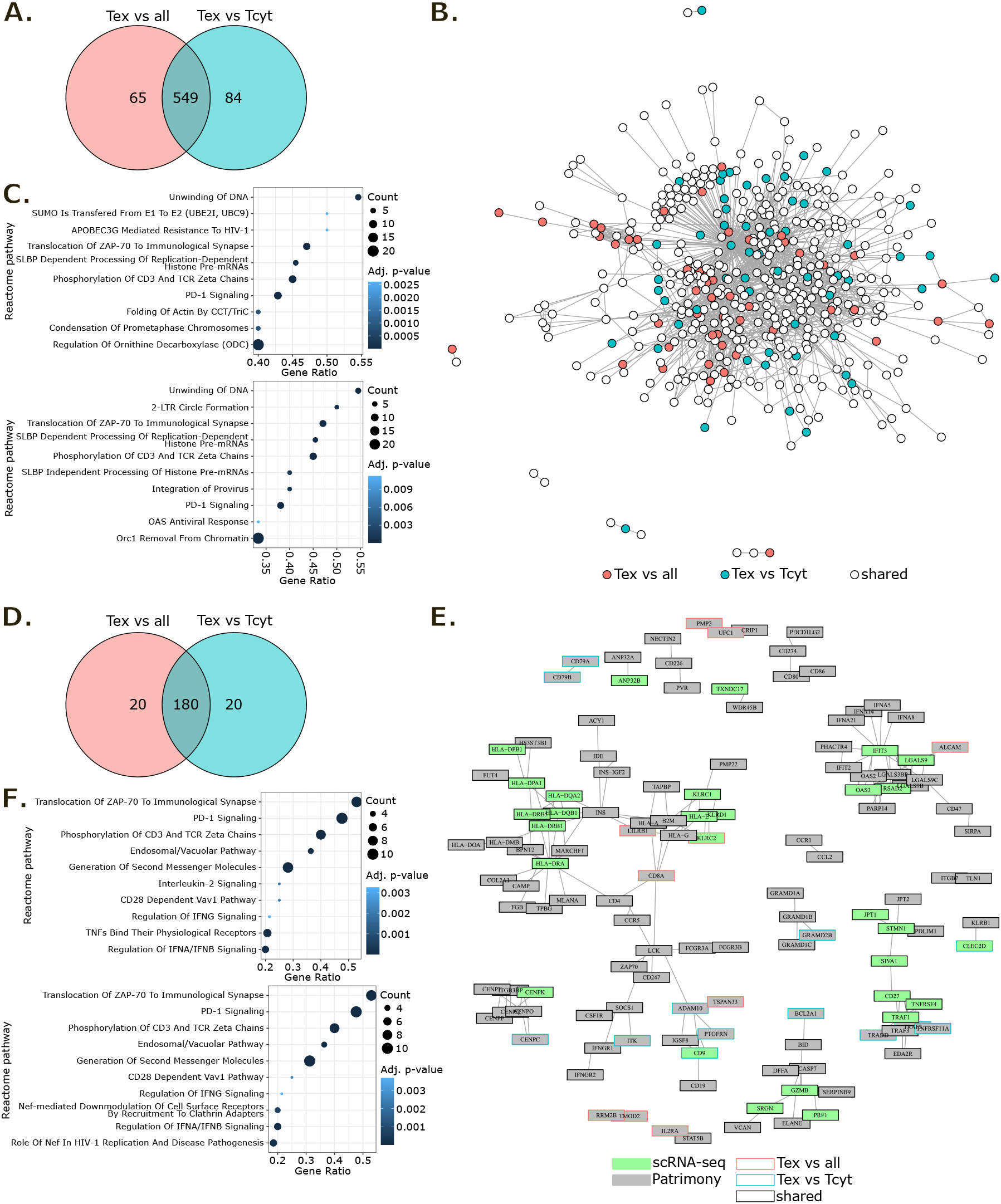
Generation of a Tex signature from the T cell atlas and its network-based rescoring for target prioritization with Patrimony. **(A)** Venn diagram of Tex scRNA-seq signatures. **(B)** Network representing direct interactions at the protein level between genes of Tex scRNA-seq signatures. Genes with no direct interactions are not represented. **(C)** Top 10 enriched Reactome pathways in “Tex vs all” (upper panel) and “Tex vs Tcyt” (lower panel) scRNA-seq gene lists. Pathways for which adjusted p-value < 0.05 and count > 2 were kept then ordered by decreasing gene ratio. **(D)** Venn diagram of Tex Patrimony top 200 genes. **(E)** Network representing direct interactions at the protein level between genes of Tex Patrimony top 200 results. Genes with no direct interactions are not represented. **(F)** Top 10 enriched Reactome pathways in “Tex vs all” (upper panel) and “Tex vs Tcyt” (lower panel) Patrimony top 200 genes. Pathways for which adjusted p-value < 0.05 and count > 2 were kept then ordered by decreasing gene ratio.

These findings provide a substantial appreciation of the transcriptomic landscape of Tex cells within the TME, highlighting pathways associated with immune response regulation, epigenetic processes and signaling cascades involved in T cell exhaustion (36, 37).

We then performed computational network analysis using the Patrimony platform on the two scRNA-seq Tex signatures. Mirroring the TI-Treg Patrimony results, the scaled diffusion scores decreased rapidly from the top 1 to approximatively the top 200 scored gene for both Tex Patrimony results (Fig S3B). Hence, we again focused on the top 200 genes for downstream analysis (Table S1). By comparing “Tex vs all” and “Tex vs Tcyt” Patrimony signatures, we observed that 90% of their top 200 genes overlapped, underscoring a higher degree of similarity than seen with the TI-Treg signatures (Fig 4D). Upon mapping Patrimony top 200 genes onto the human PPI network, several gene clusters emerged without specificities to either Tex signature (Fig 4E). We identified clusters related to antigen presentation and activation (e.g., HLA-DRA, HLA-DRB1, MLANA, CD8A, ZAP70, CD247, ZAP70, IFNGR1, etc.), cytotoxic effector functions and survival (e.g., GZMB, PRF1, SERPINB9, DEFA, BCL2A1, etc.), mitotic progression (e.g., CENPK, CENPO, etc.), TNF signaling (e.g., TNFRSF4, TRAF1, CD27, etc.), lipid metabolism and transport (e.g., GRAMD1B, GRAMD1C, GRAMD1A, GRAMD2B) and a cluster of interferon-inducible genes (e.g., IFNA8, IFNA5, IFNA14, IFNA21, OAS2, IFIT2, etc.). Interestingly, this network representation of direct interactions highlighted molecular mechanisms associated to typical persistent activation and metabolic adaptations of Tex (38, 39). While the classical PD-1-related exhaustion pathways were still evident, they did not form a largely predominant cluster.

We also conducted pathway enrichment analysis on each Tex Patrimony top 200 gene list (Fig 4F). Most of the top 10 enriched Reactome pathways in these Tex signatures were shared and related to T-cell activation. This included TCR activation, co-stimulation/repression via CD28 and PD-1, and immune regulation through downstream interferon signaling pathways. Consequently, the pathway analysis highlighted crucial aspects of CD8^+^ T cells “functional impairment”, characterized by high expression of inhibitory receptors (40), persistent antigen exposure and TCR stimulation (38).

Overall, our methodology uncovers specific modulatory mechanisms in CD8^+^ Tex cells, suggesting potential targets to counteract T cell exhaustion within the TME.

### Network propagation refined single cell gene signatures

To evaluate the advantages of applying computational network analysis on scRNA-seq gene signatures for identifying markers, we conducted a quantitative comparison between the gene signatures before and after applying the Patrimony computational method. Our analysis encompassed four scRNA-seq gene signatures (“Treg vs all”, “Treg vs Thelper”, “Tex vs all” and “Tex vs Tcyt”) each matched with four corresponding Patrimony gene signatures. We observed that in all 4 cases the number of pathways represented by at least one gene in the signatures drastically decreased after Patrimony rescoring (Fig S3C-F). This reduction was closely associated with a decrease in the number of genes within the signatures. Indeed, for “Treg vs all” as an example, a 3-fold decrease in the genes count of Patrimony signature compared to the scRNA-seq signature (Fig 3A; 3D) resulted in a 2-fold decrease in the number of represented pathways (Fig S3C). We then separately compared gene ratios and odd ratios for each represented pathway. From the gene ratio analysis, significantly represented pathways (with adjusted p-value < 0.05) were distributed on both sides of the first bisector for “Treg vs all”, “Tex vs all” and “Tex vs Tcyt” and we could observe a large number of pathways that had low to high gene ratios in the scRNA-seq signature but were no longer represented in the Patrimony signature (Fig S3C-F). For “Treg vs Thelper” which had more similar numbers of genes in the scRNA-seq and Patrimony signatures, gene ratios of significantly represented pathways were globally higher in the Patrimony signature. From the odd ratio analysis that considers the size of signatures, Patrimony signatures showed greater enrichments for a large majority of highly represented pathways in all 4 cases (Fig 5A-D). This indicates that network propagation effectively refined single cell gene signatures.

**Fig 5.**
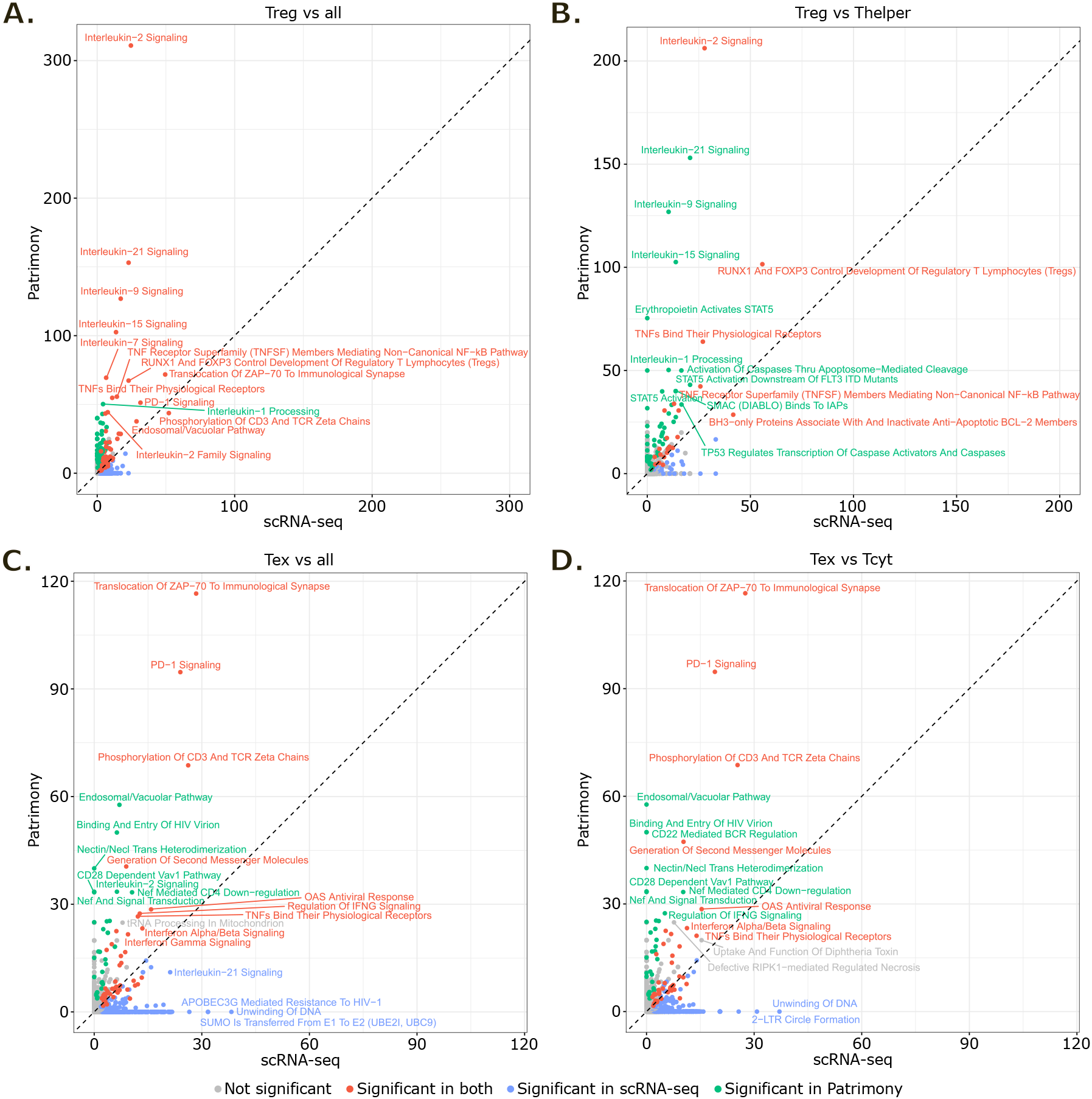
Evaluation of network propagation benefits for target prioritization. **(A-D)** Concordance plots comparing Patrimony and scRNA-seq signatures for “Treg vs all” (A), “Treg vs Thelper” (B), “Tex vs all” (C) and “Tex vs Tcyt” (D). Values correspond to odd ratios for each represented pathway. The black dashed line represents the first bisector (y=x). Pathways with adjusted p-value < 0.05 were considered significant.

In our study, we emphasized the significance of the enriched pathways identified from gene signatures after applying network propagation, underscoring their relevance in the context of existing literature. For comparison purposes, pathways analyses were performed with Fisher exact tests, thus not taking into account differential gene expression level or diffusion score. We explored the differential expression level of genes contributing to pathways highly represented both before and after network propagation or only in the scRNA-seq signatures. As an example, in the “Tex vs all” and “Tex vs Tcyt” scRNA-seq signatures, the “Unwinding of DNA” pathway was represented by 6 out of 11 genes, none of them being ranked in the top 50 genes of either signature according to their differential expression levels (log2 fold change). As for the “Translocation of ZAP-70 to immunological synapse” pathway, it was represented by 8 out of 17 genes, of which 5 were ranked in the top 50 genes of both “Tex vs all” and “Tex vs Tcyt” scRNA-seq signatures. Despite both pathways exhibiting similarly high gene ratios in the scRNA-seq signatures, their associated weight for network propagation (*i.e*. their differential gene expression levels) drove Patrimony’s resulting signatures and can explain the major enrichment difference between these pathways after network propagation. These findings confirm that network propagation effectively refined single cell gene signatures selecting genes with high biological connectivity to the weighted input signatures.

## Discussion

Targeting the immune system has shown to be a successful therapeutic approach in cancer with the approval of immune checkpoint inhibitor treatments for many tumor types. As this strategy does not act directly on malignant cells, target identification ensuring specificity and efficiency in immuno-oncology remains challenging. scRNA-seq offers the opportunity to deeply characterize any cell population of interest, opening up immuno-oncology to a wide range of possibilities. However, scRNA-seq also faces limitations such as noise and data sparsity. Here, we evaluated a methodology based on the combined use of scRNA-seq and network propagation for exploring T cell signatures and associated pathway dependencies that could eventually lead to innovative therapeutic strategies in immuno-oncology.

Two extensively studied cell populations in immuno-oncology are the CD4^+^ Treg and CD8^+^ Tex cells as their immunosuppressive functions largely contribute to cancer cells escape from immunosurveillance. Based on a new scRNA-seq pan-cancer T cell atlas, we generated gene signatures of these Tumor-Infiltrating T cell subsets. These scRNA-seq signatures were highly relevant as they were particularly enriched in genes involved in key molecular mechanisms of these populations. They shed light on processes characteristic of Tregs development through FOXP3, their survival and proliferation as well as their immunosuppressive functions through IL2 and TNF signaling. As for Tex, they featured typical exhaustion traits such as persistent antigen exposure, TCR stimulation and co-inhibitory receptors activation.

However, these scRNA-seq signatures remained very large and contained a substantial number of poorly interesting genes regarding target discovery. We thus applied computational network propagation on these signatures to rescore genes based on their biological connectivity. This approach did not fundamentally change the type of represented pathways but clearly refined the signatures selecting genes with high biological connectivity to the weighted input signatures. Furthermore, as network propagation rescores genes based on biological connectivity to the weighted input signatures, it may put forward lowly expressed or detected genes in single cell transcriptomics but for which variations could have major consequences. Such genes would then represent highly interesting therapeutic candidate targets.

The refining of gene signatures by network propagation revealed an enrichment in the endosomal/vacuolar trafficking pathway in TI-Tregs compared to other CD4^+^ T cells. Such findings align with the understanding that Tregs critically rely on oxidative phosphorylation (OXPHOS) for their energy requirements, characterized by heightened mTOR activity and moderated glycolysis levels (41). The regulation of these metabolic processes is intricately linked to the endolysosomal system, which, when impaired, has been observed to compromise Treg’s suppressive capabilities, potentially shifting their phenotype towards a more effector-like state. This shift can disrupt T cell equilibrium, inadvertently promoting antitumor responses (35). Moreover, the application of network propagation to scRNA-seq data has further delineated an augmentation in lipid metabolism and transport, particularly in Tex. This is significant considering that tumor-infiltrating CD8^+^ T cells modulate their metabolic pathways, notably through the upregulation of fatty acid oxidation, to sustain their cytotoxic function (42). Conversely, the accumulation of lipids within the TME has been implicated in the induction of CD8^+^ T cell exhaustion, facilitated by mechanisms such as increased oxidative stress (43–45, 39). These insights from network propagation analyses offer a nuanced understanding of the metabolic dependencies and vulnerabilities of T cells in the TME, highlighting potential targets for therapeutic intervention in immuno-oncology.

In conclusion, we have demonstrated that the combined use of single cell transcriptomics and network propagation could represent an innovative and effective methodology for the characterization of cell populations of interest and eventually target identification in immuno-oncology.

## Acknowledgments

The authors would like to thank all members of Servier R&D who supported the initiation and implementation of Patrimony.

## Figures description

**Fig S1. scRNA-seq T cell atlas annotation markers. (A)** UMAP representation of the T cell atlas colored by CD4^+^ or CD8^+^ subtype. **(B, C)** Bubble plot showing expression of representative markers genes of CD4^+^ (B) and CD8^+^ (C) T cell phenotypes. Color represents the normalized expression level and size represents the expression frequency. Prolif: proliferation, OXPHOS: oxidative phosphorylation, ISG: interferon-stimulated genes.

**Fig S2. Detailed cell composition analysis in the scRNA-seq T cell atlas and single cell gene signatures robustness across datasets of the atlas. (A, B)** Boxplot of CD4^+^ (A) and CD8^+^ (B) T cells detailed phenotype proportions compared to total CD4^+^ or CD8^+^ T cells in blood, adjacent normal tissue and tumor tissue. **(C)** UMAP representation of single cell Treg gene signatures scores in intra-tumor CD4^+^ T cells of the atlas. **(D)** UMAP representation of single cell Tex gene signatures scores in intra-tumor CD8^+^ T cells of the atlas. For (C) and (D), gene signature scores were computed using the *AddModuleScore* function from Seurat package with default parameters.

**Fig S3. Selection of genes for Patrimony signatures and comparison with scRNA-seq. (A, B)** Density plot of scaled diffusion scores computed with Patrimony for TI-Treg (A) and Tex (B) signatures. Vertical dashed lines correspond to the top 200 gene value for indicated signatures. **(C-F)** Concordance plots and Venn diagrams comparing Patrimony and scRNA-seq signatures for “Treg vs all” (C), “Treg vs Thelper” (D), “Tex vs all” (E) and “Tex vs Tcyt” (F). Venn diagrams show represented and significantly (sig.) represented pathways. Pathways with adjusted p-value < 0.05 were considered significant. Concordance plots compare gene ratios for each represented pathway. The black dashed line represents the first bisector (y=x).

**Table S1. scRNA-seq and Patrimony gene signatures**.

## References

1. Tang F, Barbacioru C, Wang Y, Nordman E, Lee C, Xu N et al. mRNA-Seq whole-transcriptome analysis of a single cell. Nat Methods 2009; 6(5):377–82.

2. Svensson V, Vento-Tormo R, Teichmann SA. Exponential scaling of single-cell RNA-seq in the past decade. Nat Protoc 2018; 13(4):599–604.

3. Efremova M, Teichmann SA. Computational methods for single-cell omics across modalities. Nat Methods 2020; 17(1):14–7.

4. Haque A, Engel J, Teichmann SA, Lönnberg T. A practical guide to single-cell RNA-sequencing for biomedical research and clinical applications. Genome Med 2017; 9(1):75.

5. AlJanahi AA, Danielsen M, Dunbar CE. An Introduction to the Analysis of Single-Cell RNA-Sequencing Data. Mol Ther Methods Clin Dev 2018; 10:189–96.

6. Patruno L, Maspero D, Craighero F, Angaroni F, Antoniotti M, Graudenzi A. A review of computational strategies for denoising and imputation of single-cell transcriptomic data. Brief Bioinform 2021; 22(4).

7. Ronen J, Akalin A. netSmooth: Network-smoothing based imputation for single cell RNA-seq. F1000Res 2018; 7:8.

8. Zand M, Ruan J. Network-Based Single-Cell RNA-Seq Data Imputation Enhances Cell Type Identification. Genes (Basel) 2020; 11(4).

9. Cowen L, Ideker T, Raphael BJ, Sharan R. Network propagation: a universal amplifier of genetic associations. Nat Rev Genet 2017; 18(9):551–62.

10. Guedj M, Swindle J, Hamon A, Hubert S, Desvaux E, Laplume J et al. Industrializing AI-powered drug discovery: lessons learned from the Patrimony computing platform. Expert Opin Drug Discov 2022; 17(8):815–24.

11. Desvaux E, Hamon A, Hubert S, Boudjeniba C, Chassagnol B, Swindle J et al. Network-based repurposing identifies anti-alarmins as drug candidates to control severe lung inflammation in COVID-19. PLoS One 2021; 16(7):e0254374.

12. Desvaux E, Aussy A, Hubert S, Keime-Guibert F, Blesius A, Soret P et al. Model-based computational precision medicine to develop combination therapies for autoimmune diseases. Expert Rev Clin Immunol 2022; 18(1):47–56.

13. Blaudin de Thé F-X, Baudier C, Andrade Pereira R, Lefebvre C, Moingeon P. Transforming drug discovery with a high-throughput AI-powered platform: A 5-year experience with Patrimony. Drug Discov Today 2023; 28(11):103772.

14. Korsunsky I, Millard N, Fan J, Slowikowski K, Zhang F, Wei K et al. Fast, sensitive and accurate integration of single-cell data with Harmony. Nat Methods 2019; 16(12):1289–96.

15. Gu Z. Complex heatmap visualization. iMeta 2022; 1(3).

16. Gu Z, Gu L, Eils R, Schlesner M, Brors B. circlize Implements and enhances circular visualization in R. Bioinformatics 2014; 30(19):2811–2.

17. Szklarczyk D, Gable AL, Nastou KC, Lyon D, Kirsch R, Pyysalo S et al. The STRING database in 2021: customizable protein-protein networks, and functional characterization of user-uploaded gene/measurement sets. Nucleic Acids Res 2021; 49(D1):D605–D612.

18. Kang J, Tang Q, He J, Le Li, Yang N, Yu S et al. RNAInter v4.0: RNA interactome repository with redefined confidence scoring system and improved accessibility. Nucleic Acids Res 2022; 50(D1):D326–D332.

19. Yi Y, Zhao Y, Huang Y, Wang D. A Brief Review of RNA-Protein Interaction Database Resources. Noncoding RNA 2017; 3(1).

20. Nitsch D, Gonçalves JP, Ojeda F, Moor B de, Moreau Y. Candidate gene prioritization by network analysis of differential expression using machine learning approaches. BMC Bioinformatics 2010; 11:460.

21. Liu C, Ma Y, Zhao J, Nussinov R, Zhang Y-C, Cheng F et al. Computational network biology: Data, models, and applications. Physics Reports 2020; 846:1–66.

22. Lancour D, Naj A, Mayeux R, Haines JL, Pericak-Vance MA, Schellenberg GD et al. One for all and all for One: Improving replication of genetic studies through network diffusion. PLoS Genet 2018; 14(4):e1007306.

23. Chen EY, Tan CM, Kou Y, Duan Q, Wang Z, Meirelles GV et al. Enrichr: interactive and collaborative HTML5 gene list enrichment analysis tool. BMC Bioinformatics 2013; 14:128.

24. Kuleshov MV, Jones MR, Rouillard AD, Fernandez NF, Duan Q, Wang Z et al. Enrichr: a comprehensive gene set enrichment analysis web server 2016 update. Nucleic Acids Res 2016; 44(W1):W90–7.

25. Xie Z, Bailey A, Kuleshov MV, Clarke DJB, Evangelista JE, Jenkins SL et al. Gene Set Knowledge Discovery with Enrichr. Curr Protoc 2021; 1(3):e90.

26. Wu TD, Madireddi S, Almeida PE de, Banchereau R, Chen Y-JJ, Chitre AS et al. Peripheral T cell expansion predicts tumour infiltration and clinical response. Nature 2020; 579(7798):274–8.

27. Zhang Q, He Y, Luo N, Patel SJ, Han Y, Gao R et al. Landscape and Dynamics of Single Immune Cells in Hepatocellular Carcinoma. Cell 2019; 179(4):829-845.e20.

28. Cillo AR, Kürten CHL, Tabib T, Qi Z, Onkar S, Wang T et al. Immune Landscape of Viral-and Carcinogen-Driven Head and Neck Cancer. Immunity 2020; 52(1):183-199.e9.

29. Steele NG, Carpenter ES, Kemp SB, Sirihorachai VR, The S, Delrosario L et al. Multimodal Mapping of the Tumor and Peripheral Blood Immune Landscape in Human Pancreatic Cancer. Nat Cancer 2020; 1(11):1097–112.

30. Borcherding N, Vishwakarma A, Voigt AP, Bellizzi A, Kaplan J, Nepple K et al. Mapping the immune environment in clear cell renal carcinoma by single-cell genomics. Commun Biol 2021; 4(1):122.

31. Truong L-H, Pauklin S. Pancreatic Cancer Microenvironment and Cellular Composition: Current Understandings and Therapeutic Approaches. Cancers (Basel) 2021; 13(19).

32. Kitoh A, Ono M, Naoe Y, Ohkura N, Yamaguchi T, Yaguchi H et al. Indispensable role of the Runx1-Cbfbeta transcription complex for in vivo-suppressive function of FoxP3+ regulatory T cells. Immunity 2009; 31(4):609–20.

33. Liu L, Cheng X, Yang H, Lian S, Jiang Y, Liang J et al. BCL-2 expression promotes immunosuppression in chronic lymphocytic leukemia by enhancing regulatory T cell differentiation and cytotoxic T cell exhaustion. Mol Cancer 2022; 21(1):59.

34. Noyes D, Bag A, Oseni S, Semidey-Hurtado J, Cen L, Sarnaik AA et al. Tumor-associated Tregs obstruct antitumor immunity by promoting T cell dysfunction and restricting clonal diversity in tumor-infiltrating CD8+ T cells. J Immunother Cancer 2022; 10(5).

35. Xiang H, Tao Y, Jiang Z, Huang X, Wang H, Cao W et al. Vps33B controls Treg cell suppressive function through inhibiting lysosomal nutrient sensing complex-mediated mTORC1 activation. Cell Rep 2022; 39(11):110943.

36. Sheppard K-A, Fitz LJ, Lee JM, Benander C, George JA, Wooters J et al. PD-1 inhibits T-cell receptor induced phosphorylation of the ZAP70/CD3zeta signalosome and downstream signaling to PKCtheta. FEBS Lett 2004; 574(1-3):37–41.

37. Hsu L-Y, Cheng DA, Chen Y, Liang H-E, Weiss A. Destabilizing the autoinhibitory conformation of Zap70 induces up-regulation of inhibitory receptors and T cell unresponsiveness. J Exp Med 2017; 214(3):833–49.

38. Zheng C, Zheng L, Yoo J-K, Guo H, Zhang Y, Guo X et al. Landscape of Infiltrating T Cells in Liver Cancer Revealed by Single-Cell Sequencing. Cell 2017; 169(7):1342-1356.e16.

39. Ma X, Bi E, Lu Y, Su P, Huang C, Liu L et al. Cholesterol Induces CD8+ T Cell Exhaustion in the Tumor Microenvironment. Cell Metab 2019; 30(1):143-156.e5.

40. Wherry EJ. T cell exhaustion. Nat Immunol 2011; 12(6):492–9.

41. Zeng H, Chi H. mTOR signaling in the differentiation and function of regulatory and effector T cells. Curr Opin Immunol 2017; 46:103–11.

42. Zhang Y, Kurupati R, Liu L, Zhou XY, Zhang G, Hudaihed A et al. Enhancing CD8+ T Cell Fatty Acid Catabolism within a Metabolically Challenging Tumor Microenvironment Increases the Efficacy of Melanoma Immunotherapy. Cancer Cell 2017; 32(3):377-391.e9.

43. Manzo T, Prentice BM, Anderson KG, Raman A, Schalck A, Codreanu GS et al. Accumulation of long-chain fatty acids in the tumor microenvironment drives dysfunction in intrapancreatic CD8+ T cells. J Exp Med 2020; 217(8).

44. Ma X, Xiao L, Liu L, Ye L, Su P, Bi E et al. CD36-mediated ferroptosis dampens intratumoral CD8+ T cell effector function and impairs their antitumor ability. Cell Metab 2021; 33(5):1001-1012.e5.

45. Xu S, Chaudhary O, Rodríguez-Morales P, Sun X, Chen D, Zappasodi R et al. Uptake of oxidized lipids by the scavenger receptor CD36 promotes lipid peroxidation and dysfunction in CD8+ T cells in tumors. Immunity 2021; 54(7):1561-1577.e7.

