## Supplementary figures and images for "The combined use of scRNA-seq and network propagation highlights key features of pan-cancer Tumor-Infiltrating T cells"

### Fig S1

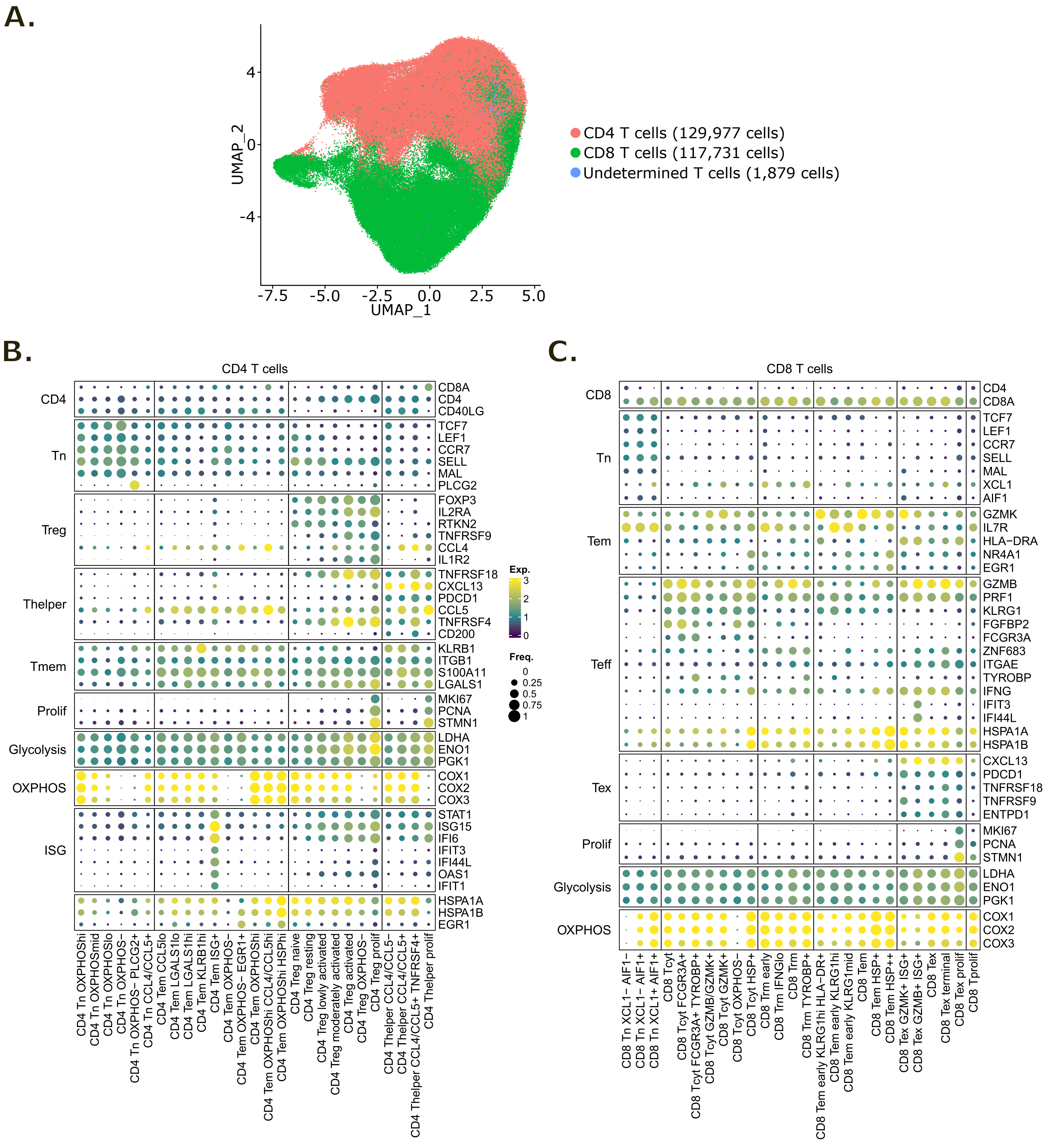

### Fig S2

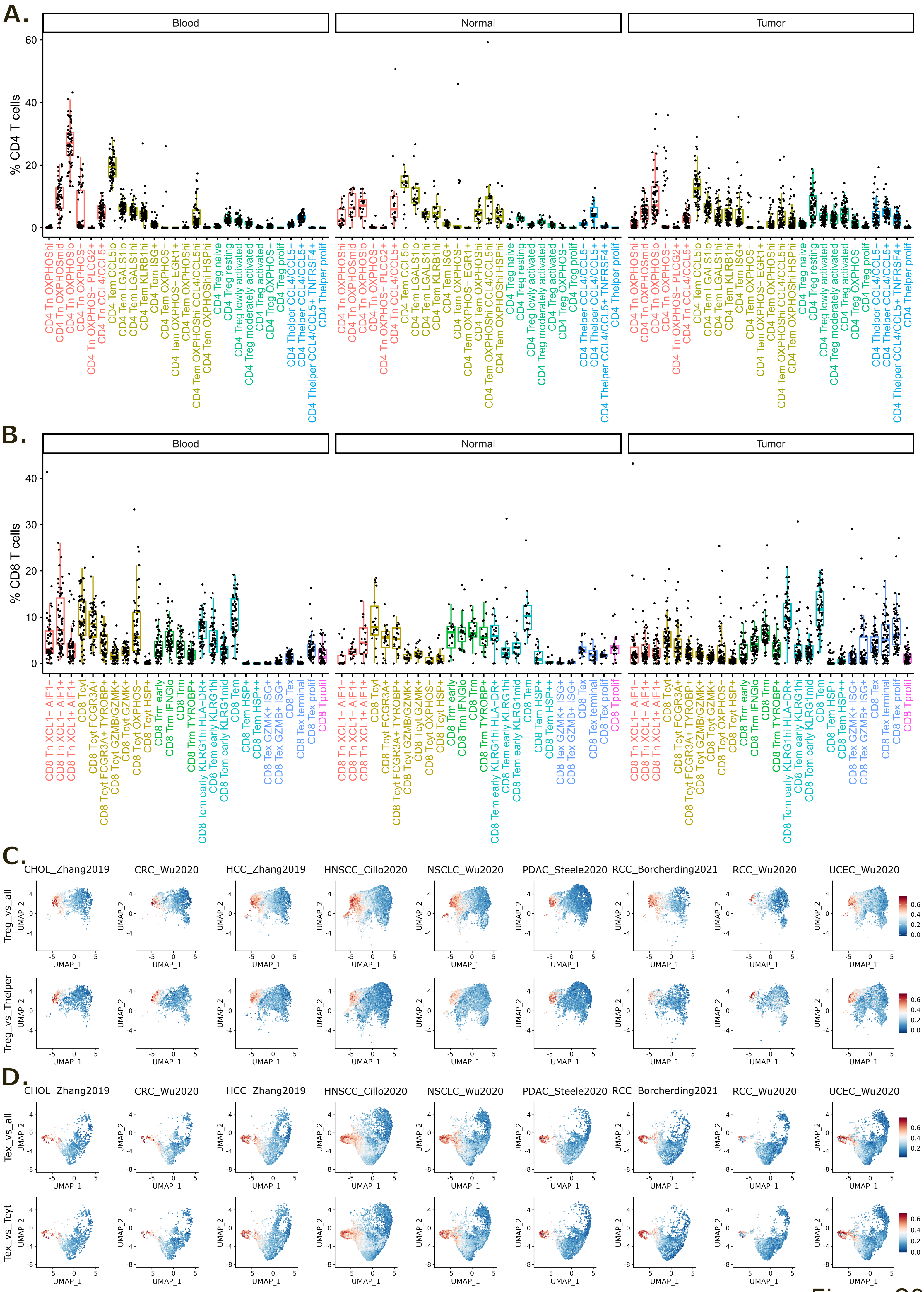

### Fig S3

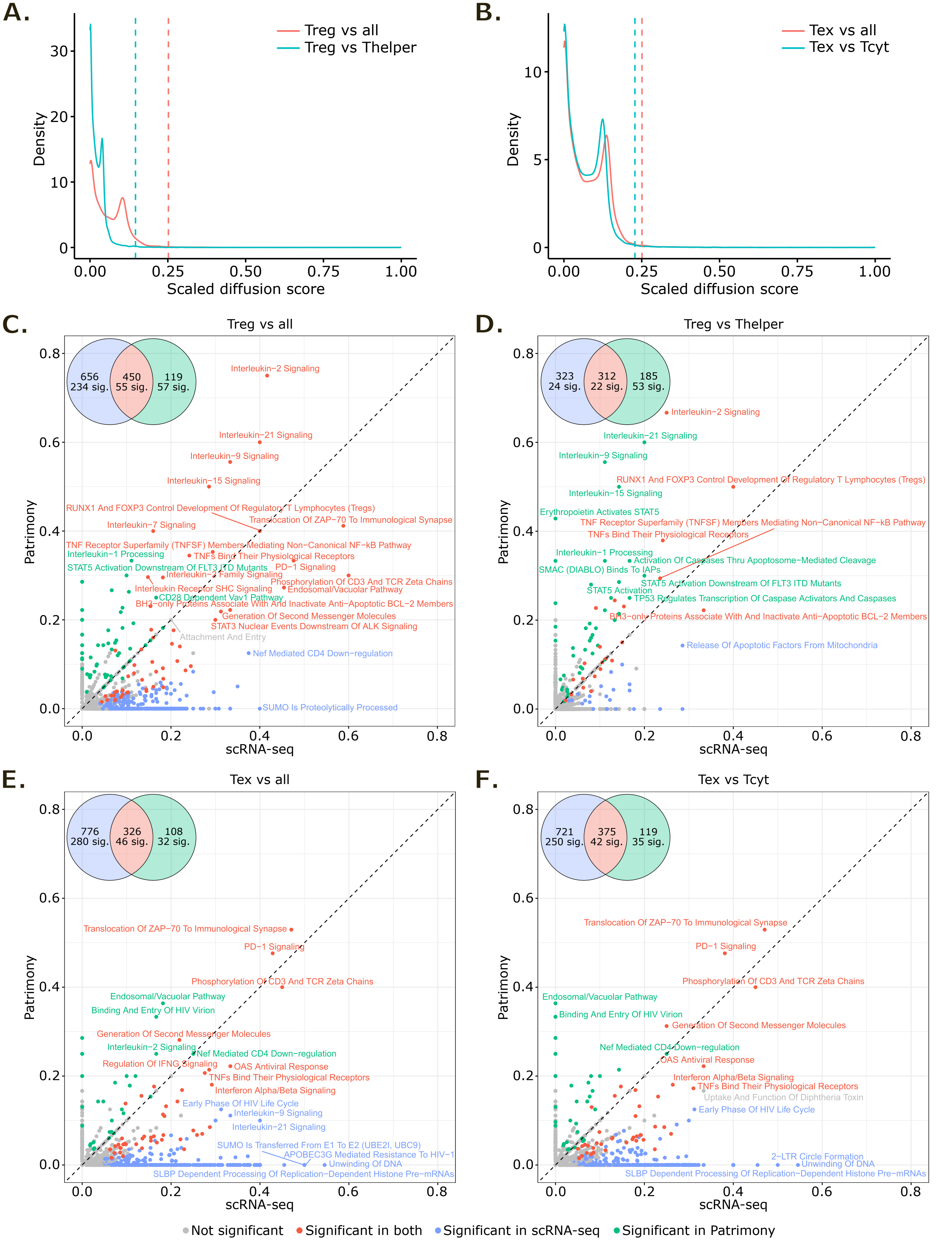
